# The mayfly subimago explained. The regulation of metamorphosis in Ephemeroptera

**DOI:** 10.1101/2021.03.17.435759

**Authors:** Orathai Kamsoi, Alba Ventos-Alfonso, Isabel Almudi, Fernando Casares, Xavier Belles

## Abstract

In the Paleozoic era, more than 400 million years ago, insects continued molting after forming functional wings. Today, however, all flying insects stop molting after metamorphosis when they become fully winged. The only exception is the mayflies (Ephemeroptera), which molt in the subimago, a flying intermediate stage between the nymph and the adult. However, the identity and homology of the subimago remains underexplored. Debate remains regarding whether this stage represents a modified nymph, an adult, or a pupa like that of butterflies. Another relevant question is why do mayflies maintain the subimago stage despite the risk of molting fragile membranous wings. These questions have intrigued numerous authors but nonetheless, clear answers have not yet been found. However, by combining morphological studies, hormonal treatments, and molecular analysis in the mayfly species *Cloeon dipterum*, we found new answers to these old questions. We observed that treatment with a juvenile hormone analog in the last nymphal instar stimulated the expression of *Kr-h1* gene and reduced that of *E93*, which suppress and trigger metamorphosis, respectively. Consequently, the subimago is not formed in these treated mayflies. This indicates that metamorphosis is determined prior to the formation of the subimago, which must therefore be considered an instar of the adult stage. We also observed that the forelegs dramatically grow between the last nymphal instar, the subimago, and the adult. This necessary growth is spread over the last two stages, which could explain, at least in part, the adaptive sense of the subimago.

## INTRODUCTION

Insect metamorphosis is the process by which an immature individual develops into a sexually reproductive adult through distinct morphological transformations. There are two metamorphosis modes: hemimetaboly and holometaboly. The hemimetabolan mode is typical of Palaeoptera, Polyneoptera, and Paraneoptera, and comprises three characteristic stages: the embryo, the juvenile instars (or nymphs), and the adult. The nymphs are morphologically similar to the adult and develop gradually until reaching the adult stage. In contrast, the Endopterygota follow the holometabolan mode, which comprises four characteristic stages, the embryo, the juvenile instars (or larvae), the pupa, and the adult. The larvae are morphologically different from the adult, and the pupa bridges the gap between these two stages (1).

In both modes, the insect stops molting after metamorphosis, when functional wings are formed (2). However, this was not the case some 400 million years ago, as shown by Paleozoic fossils of insects belonging to different orders that continued molting after acquiring functional wings (3, 4). Because membranous wings are fragile structures, shedding off the exuvia must have been a weak point in the molting process. Thus, stopping molting after forming flying wings in metamorphosis might have been favored by selective pressure, with the consequence that this became a general trait in all extant insects, except for the order Ephemeroptera, or the mayflies.

In mayflies, the molt of the last nymphal instar gives rise to a fully winged stage called the subimago, which molts again into the adult. Moreover, mayfly nymphs are aquatic and exhibit the typical adaptations to a life in water such as breathing through tracheated gills (5) and, generally, adopting herbivore-detritivore habits. Conversely, subimagos and adults are terrestrial and thus, use tracheae open through spiracles for respiration, do not usually feed, and are short-lived. Furthermore, the respective life of the subimago and the adult is very short, between a few hours and some days or a few weeks at most (6). The peculiar life cycle of mayflies raises a series of relevant questions, the most intriguing of which is about the identity and adaptive sense of the subimago. It remains unknown whether the subimago is a modified nymph, an adult, or a pupa equivalent to those of the holometabolans. In addition, it is not clear what the adaptive sense of the subimago may be.

Regarding the identity of the subimago, on the basis of morphological data, it is generally considered a kind of subadult (7, 8). However, Maiorana (9) equates the role of the subimago to that of the holometabolan pupa. Another way to assess the identity and homology of the subimago is to determine when and how the metamorphosis is triggered. Metamorphosis is regulated by two hormones: the juvenile hormone (JH), which represses metamorphosis, and 20-hydroxyecdysone (20E), which promotes the successive molts including the metamorphic one (1). In turn, the transduction mechanisms of the hormonal signals include the transcription factors Krüppel homolog 1 (Kr-h1) and E93, which are JH- and 20E-dependent, respectively. Kr-h1 plays an anti-metamorphic role, while E93 promotes metamorphosis. The interaction of both these transcription factors in the MEKRE93 pathway (10) determines whether or not metamorphosis will occur.

The MEKRE93 pathway has been described in detail in most insect orders (1), but not in mayflies. However, to properly address the above questions, this gap must be filled. The adaptive sense of the subimago is also controversial topic. Opinions vary; for example, Snodgrass (11) and Schaefer (12) argue that the subimago is a relict with no adaptive sense at all. In contrast, Maiorana (9), among other authors, contends that the subimago is necessary to complete the growth of adult structures, while Ide (13) and Edmunds and McCafferty (8) propose that, given its hydrofuge properties, the subimago facilitates the habitat transition from water to land.

In addition to morphological studies, knowledge of the mechanisms regulating metamorphosis would also help shed some light on the issues discussed above. However, practically nothing is known about these mechanisms in paleopterans, including mayflies. Thus, our approach was to elucidate them at molecular level, with special emphasis on those involved in the formation of the subimago, and to examine how these mechanisms compare to those described in neopteran insects. We used the ovoviviparous species *Cloeon dipterum* as a model because an efficient system for breeding it in the laboratory has been established (14), and because its genome has been sequenced and published, which provides a solid base of genetic information (5). *C. dipterum* exhibits the longer, ancestral pattern of longevity (8), which includes a nymphal period of about 30–50 days that develops through 13–19 instars, the subimaginal stage that lasts about 1 day, and the adult that can live 10–20 days, which is exceptionally long for mayflies (14).

In this work, our molecular studies showed that metamorphosis is hormonally determined prior to the formation of the subimago, which must therefore be considered an instar of the adult stage. Moreover, our morphological data indicate that forelegs dramatically grow between the last nymphal instar, the subimago and adult. This necessary growth is spread over the last two stages, which could explain, at least in part, the adaptive sense of the subimago.

## RESULTS

### The wings mature during the transition from the last nymphal instar to the subimago and adult

To characterize the developmental stages preceding metamorphosis in *C. dipterum*, we examined the last four nymphal instars, using the wing pads as a diagnostic feature. The nymphal instars examined were the pre-antepenultimate (PAN), antepenultimate (AN), penultimate (PN), and last (LN). *C. dipterum* only has one pair of wings that are in the mesothorax; thus, we used the mesothoracic wing pad length to identify the above-mentioned nymphal instars. In the PAN, the wing pads do not reach the first abdominal segment (A1) (Fig. 1A). In the AN, the wing pads just reach the anterior edge of A1 (Fig. 1B). In the PN, the wing pads do not reach the second abdominal segment (A2) (Fig. 1C). Finally, in the LN, wing pads are much longer, going beyond A2 (Fig. 1D).

**Fig. 1.**
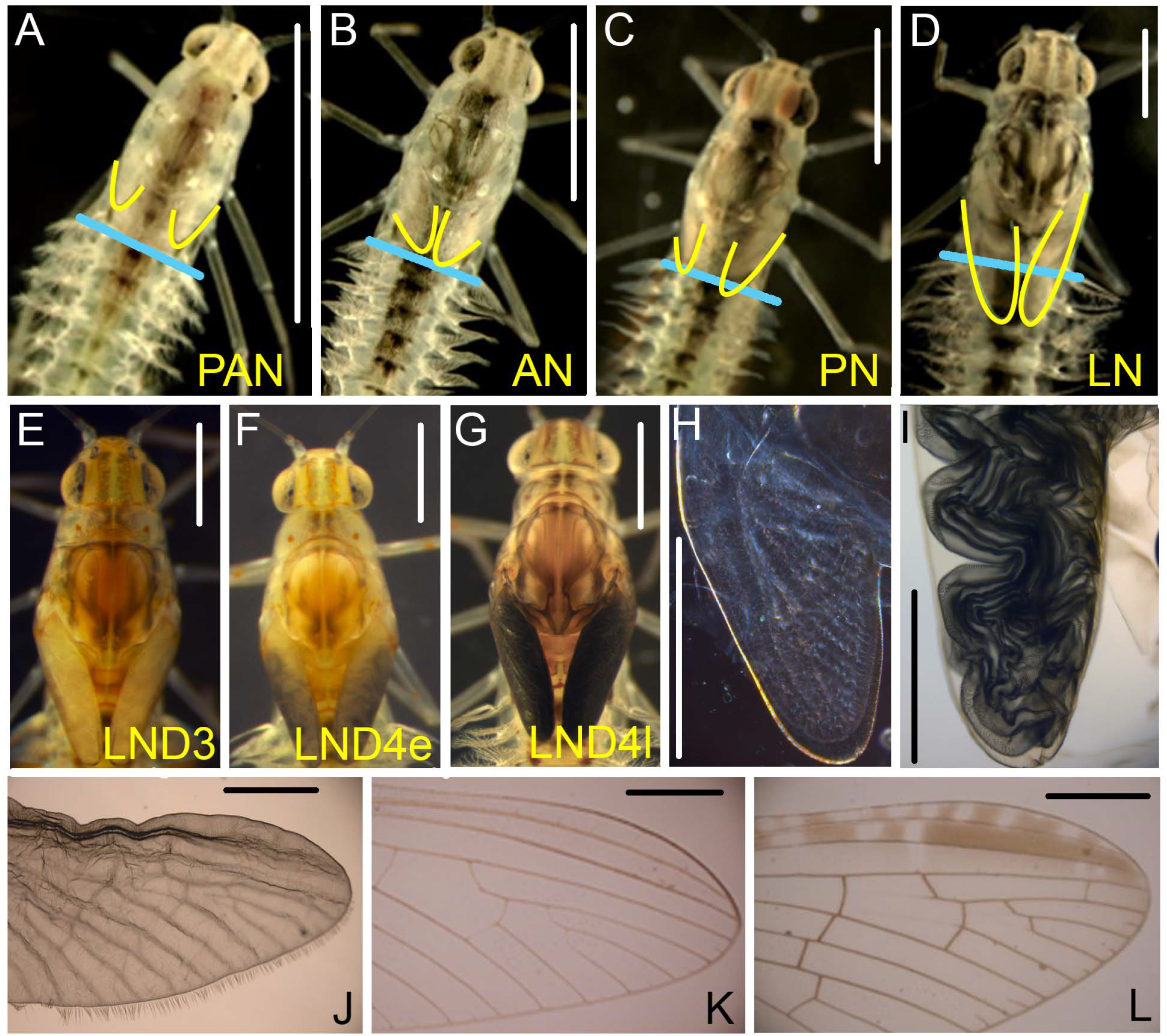
Last nymphal instars and wing maturation in *Cloeon dipterum*. (*A*) Female nymph in the pre-antepenultimate instar (PAN); the wing pads do not reach the first abdominal segment (A1). (*B*) Female nymph in the antepenultimate instar (AN); the wing pads just reach the anterior edge of A1. (*C*) Male nymph in the penultimate instar (PN); the wing pads go beyond the anterior edge of A1. (*D*) Female nymph in the last instar (LN); the wing pads go beyond A2. (*E-G*) Development of the wing pads in the LN; shape and color in 3-day-old nymphs (LND3) (*E*), early 4-day-old (LND4e) (*F*), and late 4-day-old (LND4l) (*G*). (*H*) Wing primordia in the PN. (*I*) Wing in LN4l, folded within the wing pad. (*J*) Wing in LND4l, artificially removed from the wing pad and extended on a slide. (*K*) Apical part of a subimago wing. (*L*) Apical part of an adult wing. From (*A*) to (*D*), images obtained on the first day of the corresponding instar; the perimeter of the wing pads has been indicated with a yellow line, and the anterior edge of A1 with a blue line. Scale bars: 1 mm.

During the LN, the wing pads gradually change color along the instar, as seen (Fig. 1 D-G) at 2 h (LND0), 24 h (LND1), 48 h (LND2), 72 h (LND3), and 84 h (LND4) after molting, after which the nymphs molt to subimago. From LND0 to LND2, the wing pads are thin, semi-transparent, and have a pale gray-yellow color (Fig. 1D). In LND3 the wing pads become thicker and the color changes to an intense yellow (Fig. 1E). In LND4, the color of the wing pads gradually changes from partially gray early on day 4 (Fig. 1F) to black at the end of the instar (Fig. 1G). The changes observed during LN, especially during LND4, suggest that the transformation into mature wings occurs towards the end of this instar. Indeed, while the wing pads of the PN contain only wing primordia (Fig. 1H), those of LND4 contain a folded wing (Fig. 1I), which can be artificially taken out of the wing pad and extended on a slide Fig. 1J). In terms of venation, coloration, and pubescence, this wing corresponds to that of a normally ecdysed subimago (Fig. 1K). Moreover, it is dull and translucent, in part because the adult wing is developing underneath that of the subimago. The definitive adult wing (Fig. 1L) is shiny and transparent, and does not have any hairy structures.

### The mouthparts are lost during metamorphosis while the legs lengthen and change shape

The mouthparts of *C. dipterum* nymphs consist of a pair of strong mandibles, a flap-like labrum, maxillae, labium, and hypopharynx. In contrast, the subimago and the adult do not feed and lack mouthparts (15). Thus, the newly formed mouthparts can be observed through the transparent exoskeleton at the end of each nymphal instar (Fig. 2A), except during the last one when the subimago is being formed, which does not have mouthparts (Fig. 2B).

**Fig. 2.**
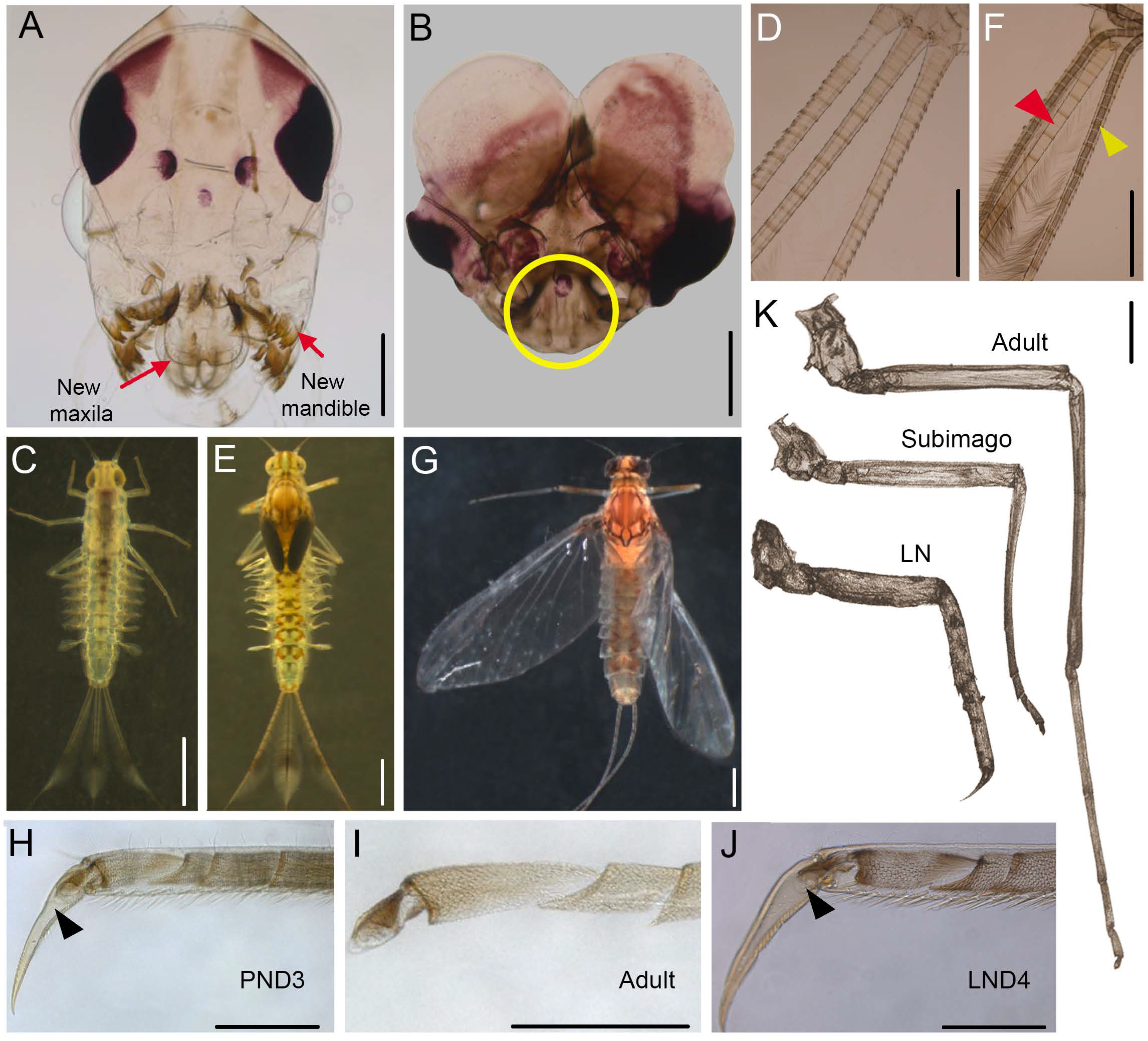
The mouthparts, central filament and legs during *Cloeon dipterum* metamorphosis. (*A*) Head of a female in the last day of the penultimate nymphal instar showing the corresponding mouthparts, and a new set of mouthparts (exemplified by a new pair of mandibles and maxillae) that are being formed, which corresponds to the last nymphal instar (LN). (*B*) Head of a male subimago showing that mouthparts disappeared (circle). (*C*) Fourth nymphal instar with two cerci and a central filament. (*D*) Detail of the cerci and filament in the fourth nymphal instar. (*E*) Late sixth nymphal instar (black wing pads stage) showing the two cerci and the central filament. (*F*) Detail of the cerci and filament in a late sixth nymphal instar after the apolysis; note the new cerci formed after the apolysis (yellow arrowhead), whereas the central filament only show the nymphal structure (red arrowhead). (*G*) Habitus of a female subimago. (*H*) Detail of the apical tarsomere in 3-day-old penultimate nymphal instar (PND3); note the actual claw and the new claw formed after the apolysis and corresponding to the LN (arrowhead). (*I*) Detail of the apical tarsomere in the adult. (*J*) Detail of the apical tarsomere in 4-day-old LN (LND4); note the small hook-shaped structure formed after the apolysis and corresponding to the subimago (arrowhead). (*K*) Foreleg of a male LN, subimago and adult. Scale bars: 0.25 mm in (*H*), (*I*), and (*J*); 0.5 mm in (*A*), (*B*), (*D*), and (*K*); 1 mm in (*C*), (*E*), (*F*), and (*G*).

After hatching, *C. dipterum* nymphs have only two caudal cerci. The central filament develops after two or three molts (Fig. 2C and D), gradually increasing in size at each molt until LN (Fig. 2E and F). However, the subimago and the adult of this species do not have a central filament (Fig. 2G). According to our observations in LND4, after the apolysis and formation of the new cuticle, new respective subimaginal cerci are formed, but not the central filament (Fig. 2F).

In the legs, the apical tarsomere is remodeled during metamorphosis. In nymphs, it is elongated and claw-shaped (Fig. 2H), whereas in the subimago and the adult it is hook-shaped (Fig. 2I). The formation of this new structure can be observed in LND4, where it can be seen through the transparent exoskeleton that the subimaginal hook-shaped structure is formed instead of the nymphal long claw (Fig. 2J). Furthermore, the forelegs dramatically grow between the LN and the adult, the length practically doubling, in both males and females (Fig. 2K).

### Metamorphosis is genetically determined in the last nymphal instar

Between the PAN and adult stage, we studied the expression of the *Krüppel homolog 1* (*Kr-h1*) and *Ecdysone-induced protein 93F* (*Eip93F* or *E93*) genes, which are both direct regulators of metamorphosis (10). We also examined *Broad complex* (*Br-C*), involved in wing development in hemimetabolans (16–18), and the 20E-dependent gene *Hormone receptor 3* (*HR3*), as a readout of 20E and a molecular marker of apolysis (19). The expression was studied in females in the mid PAN, AN, and PN, on every day of the LN, in freshly emerged subimagos, and in one-day-old adults.

The results indicate that *Kr-h1* expression abruptly decreases just after molting to LN (LND0) and that these values remain low until LND3; expression then dramatically increases during LND4 and subimago, remaining relatively high in the adult (Fig. 3). In parallel, *E93* expression is extremely low before LN, starts increasing during LND0, stays relatively stable from LND1 to LND3, and then notably increases during LND4, maintaining similarly high values in the subimago and adult (Fig. 3). The inverse expression patterns of *Kr-h1* and *E93* between PN and LND3 are expected, since Kr-h1 represses *E93* (10), but parallel high expression values of *Kr-h1* and *E93* in LND4, the subimago, and adult is intriguing, suggesting that the two genes are expressed in different tissues at different levels in these stages. *HR3* expression peaks in LND3 and in the subimago, indicating that there are discrete 20E pulses during these stages, triggering respective molting processes.

**Fig. 3.**
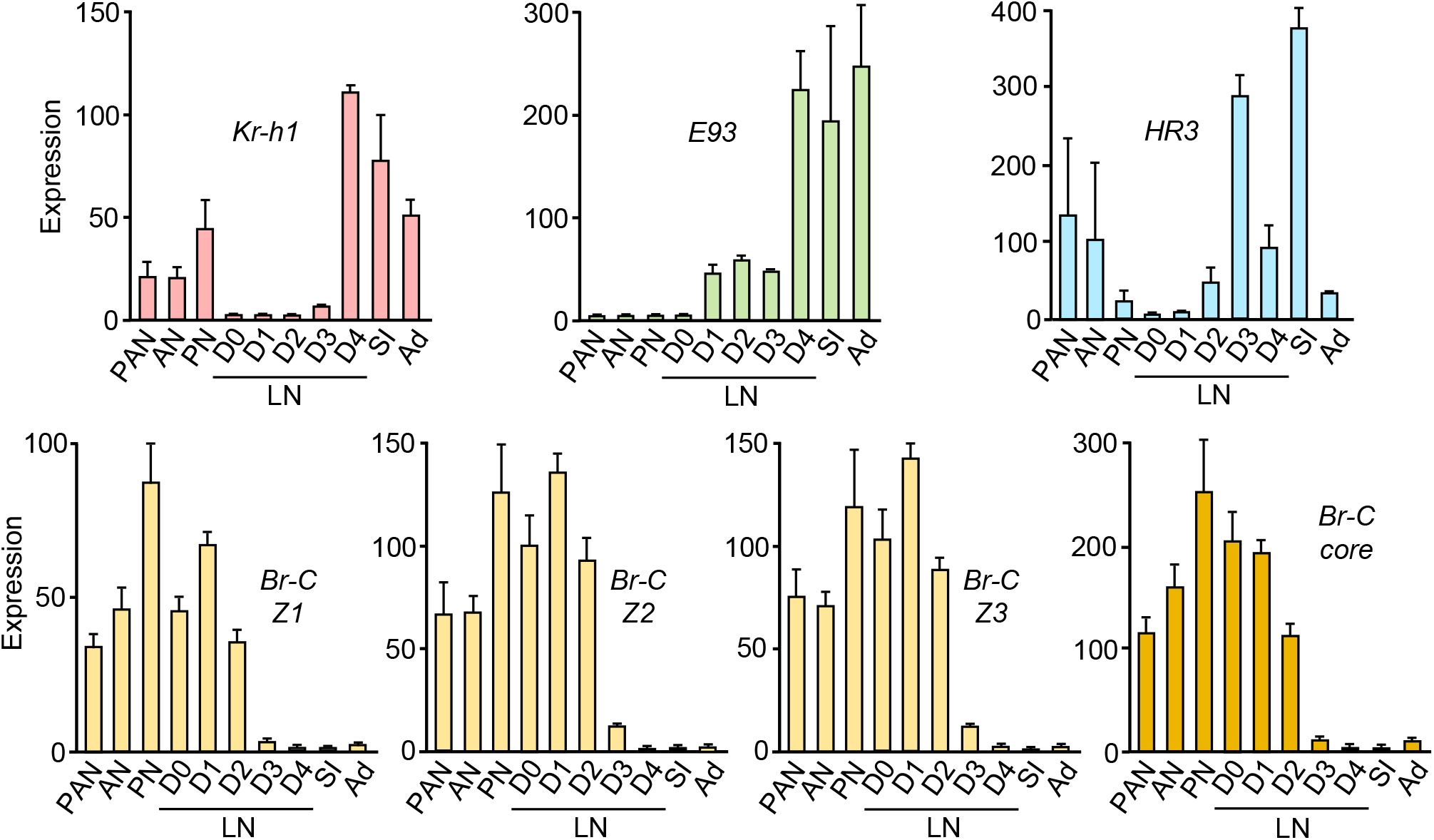
Expression of genes relevant for metamorphosis in *Cloeon dipterum*. The genes *Krüppel homolog 1* (*Kr-h1*), *Ecdysone-induced protein 93F* (*Eip93F* or *E93*), *Hormone receptor 3* (*HR3*) and *Broad-complex* (*Br-C*) were examined. In the latter gene, we measured the expression of each of the isoforms, *Br-C Z1, Br-C Z2* and *Br-CZ3*, and the joint expression of all the isoforms (*Br-C core*). The expression was measured in females of pre-antepenultimate nymphal instar (PAN), antepenultimate nymphal instar (AN), penultimate nymphal instar (PN), all days of the last nymphal instar (LN, from D0 to D4), the subimago (SI) and the adult (Ad). The results are indicated as copies of the examined mRNA per 1000 copies of CdActin-5c mRNA, and are expressed as the mean ± SEM (n=3).

Finally, we examined the *Br-C* gene, in which genomic analyses revealed the occurrence of three isoforms associated with zinc fingers, which are homologous to Z1, Z2 and Z3 of other insects (*SI Appendix*, Figs. S1-S2). Therefore, we measured the expression of each of the isoforms (Br-C Z1, Br-C Z2 and Br-C Z3) with specific primers designed to distinguish the zinc finger regions, as well as the global expression of all the isoforms with primers designed to recognize the region common to all of them (Br-C core). The expression of all three Br-C isoforms followed a similar pattern: it increased up to PN, maintained relatively high expression levels until LND2, fell in LND3, and then maintained very low levels until the adult stage. *Br-C Z2* and *Br-C Z3* were expressed at similar levels, whereas *Br-C Z1* expression was just over half the level of *Br-C Z2* and *Br-C Z3* (Fig. 3).

### Treatment with a juvenile hormone mimic inhibits metamorphosis

We administered the JH mimic methoprene during LN to assess whether it could prevent metamorphosis in *C. dipterum*, as occurs in neopteran insects. Treatments with doses of 0.1 µg, 1 µg, or 5 µg had no effect when topically applied on freshly ecdysed LN; only a dose 50 µg of methoprene produced an inhibitory effect on metamorphosis. Control insects (*n* = 28) molted normally to subimago and then to adults. In contrast, methoprene-treated insects (*n* = 25) arrested on the last day of the instar and died one or two days later.

Examination of these methoprene-treated insects in late LND4 revealed that they had completed the formation of a new cuticle, but that this corresponded to a supernumerary nymph rather than a subimago. Regarding the head, the controls did not form mouthparts, as expected when molting to a subimago, whereas a new set of mouthparts was developed in methoprene-treated insects, which would normally correspond to nymphal development (Fig. 4A). Moreover, the controls formed new cerci, but they did not form the central filament, in line with the morphology of the subimago, whereas methoprene-treated insects formed two cerci and the central filament, corresponding to a nymph-to-nymph transition (Fig. 4B).

**Fig. 4.**
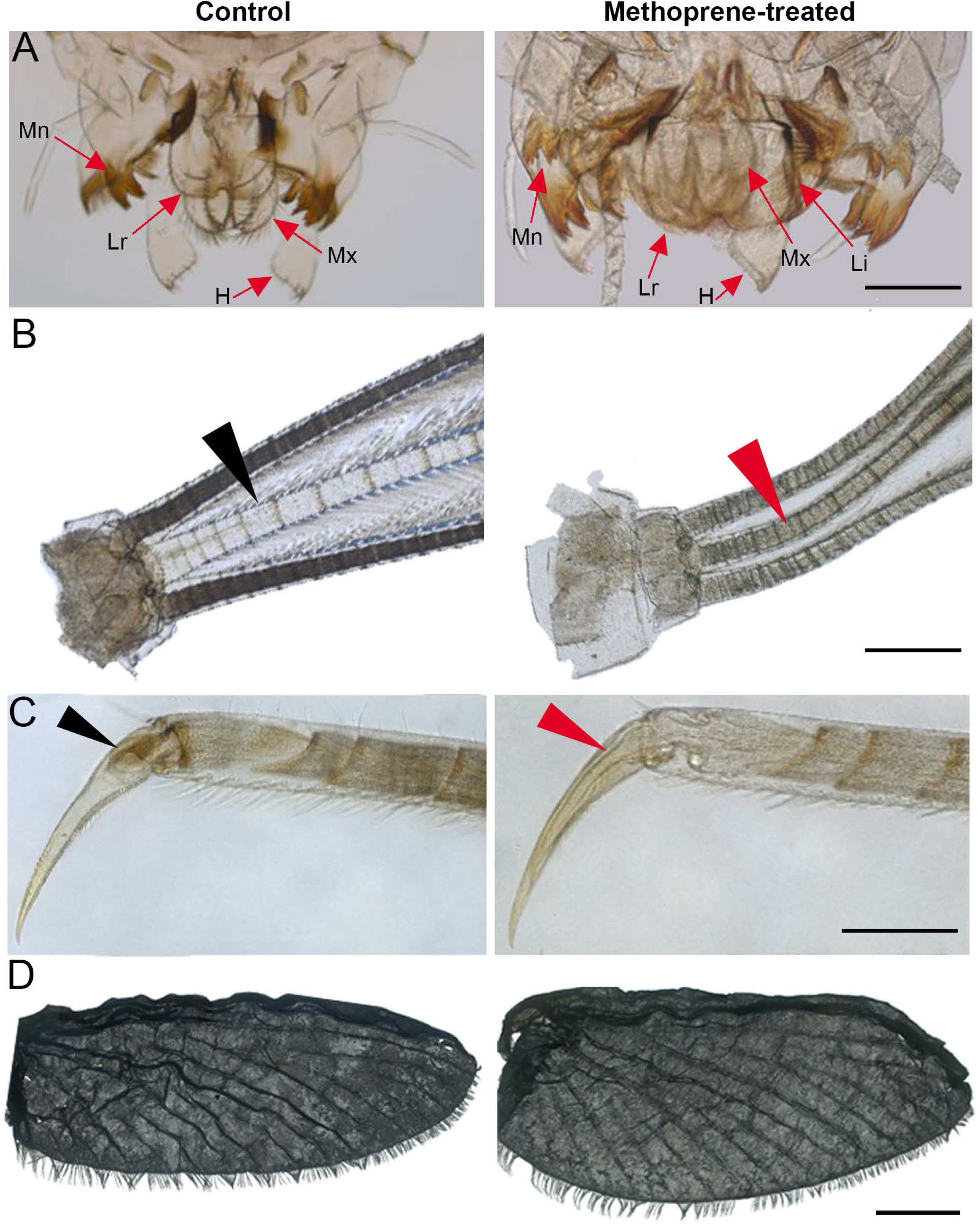
Effects of methoprene on metamorphosis in *Cloeon dipterum*. Methoprene was administered in freshly emerged last instar nymphs (LND0), and morphological features were examined four days later (LND4) in controls and methoprene-treated insects. (*A*) Mouthparts; note that the controls only show the nymphal structures, whereas the methoprene-treated show the nymphal structures and a new set formed after the apolysis; H: hypopharynx, Li: labium; Lr: labrum; Mn: mandible; Mx: maxilla. (*B*) Cerci and central filament; note that the controls form new cerci but they do not form the central filament (black arrowhead), whereas the methoprene-treated insects form the cerci and the central filament (red arrowhead). (*C*) Last tarsomere exemplified by the hind leg; note that the controls form the new structure, short and hook-shaped, which is characteristic of the subimago (black arrowhead), whereas the methoprene-treated insects form again the elongated and claw-shaped structure that is characteristic of the nymphal instars (red arrowhead). (*D*) Developing wings; the general morphology and vein pattern are similar in controls and in methoprene-treated. Scale bars: 0.5 mm in (*A*) and (*C*); 1 mm in (*B*), and 2 mm in (*D*).

With respect to the last tarsomere, controls developed the hook-shaped structure characteristic of the subimago, whereas the methoprene-treated insects again formed the same claw-shaped structure of the nymphal instars (Fig. 4C). Regarding the wing pads, after dissecting out the developing wings and extending them on a slide, a similarly shaped wing was observed in both the controls and the methoprene-treated insects. In both cases, the wing morphology, vein patterning and row of cilia on the edges, corresponded to a subimago wing (Fig. 4D).

### Treatment with a juvenile hormone mimic modifies the expression of metamorphosis genes

Two days after treatment with methoprene, we measured the expression of *Kr-h1, E93* and *Br-C*. We quantified the mRNA levels in the head (which, in addition to nervous tissues and the eye system, contains the mouthparts), wing pads (the wing primordia and pterothecae that protect them), abdominal epidermis (the carcass remaining after removing all the abdominal content), and the ovaries. The results (Fig. 5) showed that, in the head and the wing pads, the expression of *Kr-h1* and all the *Br-C* isoforms had increased while *E93* expression had significantly decreased in the insects treated with methoprene. The results for the abdominal epidermis were similar, although the tendency towards decreased *E93* expression was not statistically significant. In contrast, methoprene treatment was not associated with significant changes in expression for any of the genes when measured in the ovaries (Fig. 5).

**Fig. 5.**
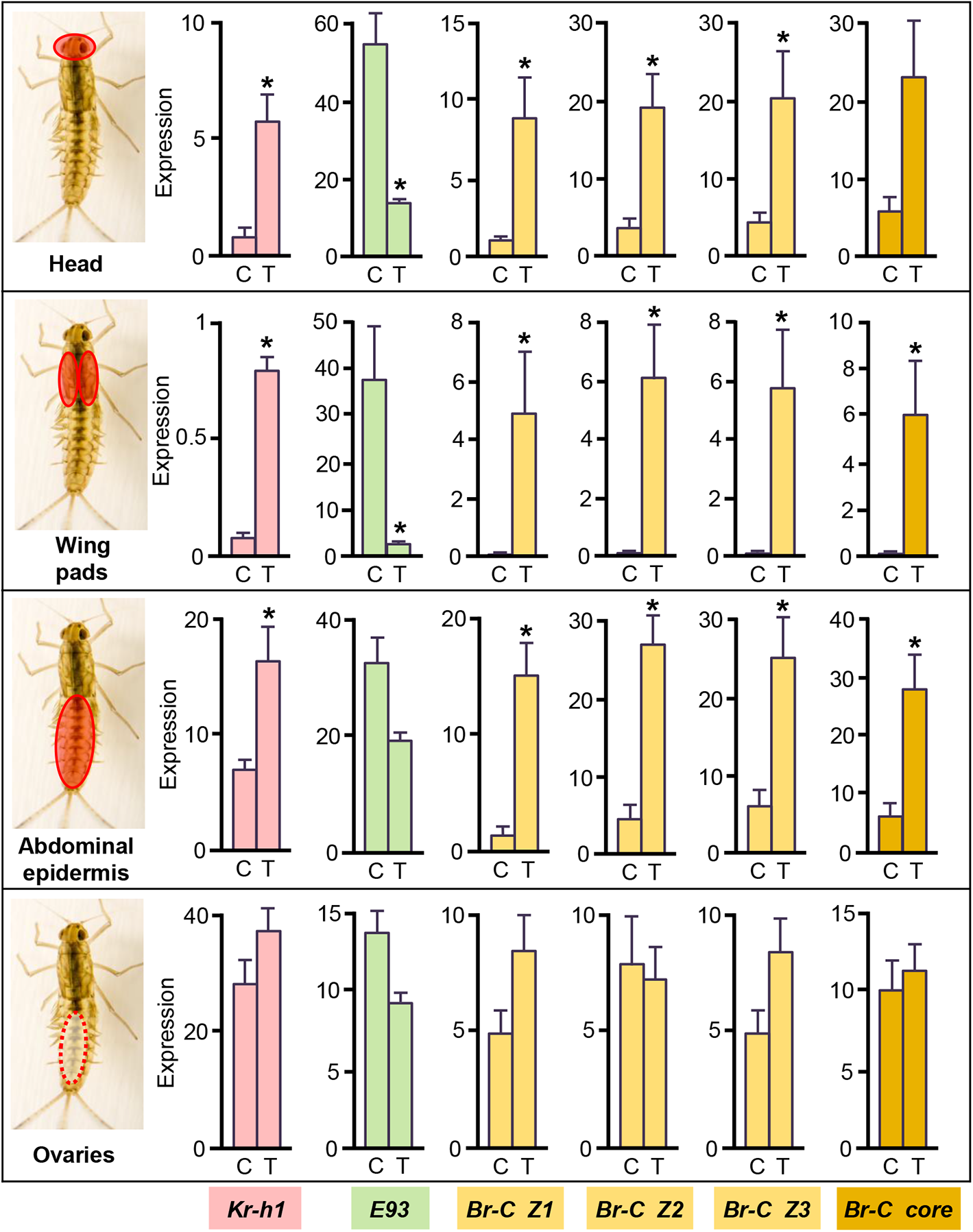
Effects of methoprene treatment on the expression of genes involved in metamorphosis in *Cloeon dipterum*. Methoprene was administered in freshly emerged last instar female nymphs, and gene expression was measured in the head, wing pads, abdominal epidermis, and ovaries two days later. The genes *Krüppel homolog 1* (*Kr-h1*), *Ecdysone-induced protein 93F* (*Eip93F* or *E93*) and *Broad-complex* (*Br-C*) were examined; in the latter gene, we measured the expression of each of the isoforms, *Br-C Z1, Br-C Z2* and *Br-C Z3*, and the joint expression of all the isoforms (*Br-C core*). The results are indicated as copies of the examined mRNA per 1000 copies of CdActin-5c mRNA, and are expressed as the mean ± SEM (n=3); the asterisk indicates statistically significant differences between methoprene-treated and control insects (p<0.05) according to the REST (31).

## DISCUSSION

### The morphological data

The nymphs of *C. dipterum* have a buccal apparatus adapted for chewing, whereas the subimago and adult, which do not feed, have no mouthparts (15). Our study revealed that the mouthparts do not form in the LN-subimago transition. Another structure that is lost during this transition is the central filament. Most mayfly species retain not only the two caudal cerci, but also the central filament in the subimago and adult stages, structures that apparently contribute to the aerodynamic stabilization of flight (20). In *C. dipterum*, however, the subimago and adult stages do not possess the central filament, which, according to our observations, is no formed during the LN–subimago transition.

In contrast, *C. dipterum* metamorphosis involves the formation of new structures. The most obvious is the pair of membranous wings developed in the mesothorax upon the formation of the subimago, with this wing pair developing again with the formation of the adult. In parallel, the last tarsomere of the leg is transformed during metamorphosis. In nymphs it is claw-shaped, which helps the insect cling to the bottom of streams. In contrast, it is hook-shaped in the subimago and the adult, which is more useful for attaching to terrestrial substrates and, in the male, for grasping the female during in-flight mating (6). The tarsomere transformation takes place in the transition from LN to subimago, whereas in the transition from subimago to adult, the hook-shaped last tarsomere is formed again.

This repeated sequential formation of the tarsus in the subimago and adult becomes contracted in species with an exceptionally short longevity pattern, in which the subimago lives only a few minutes. In *Palingenia fuliginosa*, for example, both the subimaginal and adult tarsi develop nearly simultaneously, and can be seen overlapping late in last instar nymphs (21). In parallel, the legs grow in length between the LN and the adult stages of *C. dipterum*, especially the tarsi of the male, which are more than two times longer in the adult than in the LN. Spectacular tarsi lengthening has also been observed in other species. For example the foreleg tarsi of the adult male of *Ephoron leukon* are between five and seven times their nymphal length (13), and that of *P. fuliginosa* is about eight times longer than in the nymph (8).

### The gene expression patterns

Regarding the molting processes, the short duration of the subimago stage might suggest that a single pulse of 20E at the end of LN would trigger the formation of the superimposed subimago and adult structures. However, the expression pattern of HR3 clearly suggests that there is one 20E pulse in LND3 and another in the subimago, which correspond to the respective molting processes to subimago and the adult. Regarding metamorphosis, the daily values during LN also indicate that the expression of *Kr-h1* dramatically decreases from the beginning of LN until LND3, suggesting that a Kr-h1-free period, which would allow an increase of *E93* expression through the MEKRE93 pathway, is crucial for metamorphosis in *C. dipterum* (10).

With respect to wing formation, *Br-C* expression steadily declines during LN, almost completely vanishing between LND3 and LND4, just when the wings become fully mature. In hemimetabolan neopteran insects, Br-C plays a crucial role in promoting wing primordium growth and development within the wing pads (16, 18, 22), and it is likely that it plays the same role in *C. dipterum*. Interestingly, *Br-C* expression is very low in the subimago, which suggests that the discrete wing transformation that occurs in the transition from subimago to adult (loss of cilia and changes in pigmentation and shine) do not require Br-C.

As regards Br-C isoforms, there seems to be a certain tendency to reduce their number through insect evolution, from early-branching insect species, such as the cockroach *Blattella germanica*, which has six (23), to the most modified species, such as the fly *Drosophila melanogaster* which has four (24). Therefore, although we only found three, we do not rule out that *C. dipterum* may have other Br-C Zn finger isoforms.

### The methoprene experiments and the regulation of metamorphosis

Treatment with methoprene in LND0 prevented the formation of the subimago, with a supernumerary nymphal instar being formed instead. These nymphs were unable to ecdyse, but their morphological characteristics observed through the transparent exoskeleton (absence of mouthparts and the presence of a central filament and claw-shaped apical tarsomere) clearly revealed their nymphal character. In contrast, the methoprene-treated insects were able to develop normally shaped and patterned wings.

At the molecular scale, methoprene treatment triggered a significant increase in *Kr-h1* mRNA levels in the head, wing pads, and abdominal epidermis. This result is not surprising since JH induces *Kr-h1* expression in metamorphosing tissues in hemimetabolan, neopteran insects (22, 25). The increase in *Kr-h1* expression was accompanied by a parallel decrease in *E93* expression, which makes sense as Kr-h1 represses *E93* expression through the MEKRE93 pathway. Indeed, the E93 deficiency explains the formation of a supernumerary nymph instead of a subimago, given that this factor triggers metamorphosis (26). We must recognize that the interaction between Kr-h1 and E93 might be best assessed through RNAi approaches. However, our attempts using distinct dsRNAs designed to specifically target Kr-h1 and E93 failed to deplete either of the two respective transcripts.

Methoprene also induced a significant increase in *Br-C* expression in metamorphosing tissues, which is consistent with the observation that JH enhances the expression of *Br-C* during the nymphal period of hemimetabolan, neopteran insects (18). In contrast with the results obtained for the head, wing pads, and abdominal epidermis, methoprene did not significantly affect the expression of *Kr-h1, E93* and *Br-C* in the ovaries. No effects in the ovary of other species have been reported for JH in terms of the expression of these factors. In any case, our results suggest that they would have been unaffected by this hormone, at least in *C. dipterum* and under the conditions of our experiments.

### The homology and the adaptive sense of the subimago

The intersection of *Kr-h1* and *E93* expression patterns in LN indicates that metamorphosis takes place in the transition from the LN to the subimago. In neopterans, this intersection occurs in the last nymphal instar (hemimetabolans) or in the pupa (holometabolans), and marks the activation of adult morphogenesis (data reviewed in (1)). Accordingly, the subimago should be considered as the first instar of the adult stage, with the “adult” being the second and final instar.

This conclusion is also supported by comparative studies of the transcriptomes of young larva, mature larva, subimago, and adult of the mayfly *Cloeon viridulum*, which showed that the transcriptome most similar to that of the subimago is that of the adult (27). Under these premises, the metamorphosis mode of mayflies is hemimetabolan, with juveniles being morphologically like adults, and without undergoing a pupal stage. Thus, other terms proposed to designate the mayfly metamorphosis, such as paurometaboly (28) or prometaboly (29), are misleading and should be discouraged.

The adaptive sense of the subimago has been explained by Ide (13) in terms of the hydrofuge properties of the hairy body, leg, and wing surfaces in this stage, which would facilitate the water–air transition immediately after metamorphosis. Edmunds and McCafferty (8) support the same argument with experiments showing the hydrofuge capabilities of the subimago of various species, which are not exhibited by the adult.

From a different perspective, Maiorana (9) proposed that the adaptive sense of the subimago is to allow for the growth necessary to transit from the nymphal to the adult morphology. This idea is generally supported by data showing that full expansion of body structures, notably legs and caudal cerci, as well as genital structures, is completed with the formation of the adult. Long cerci contribute to flight stabilization (20), whereas long legs terminating in a hook, while also contributing to flight optimization, are also required in males for grasping the female during mating (6). Our observations on the growth of the fore leg, which is distributed in the LN–subimago and subimago–adult transitions, concur with the proposal by Maiorana (9).

Natural selection operates upon the products of chance, but in a domain of demanding conditions, necessity prevails (30). The dramatic shortening of the terrestrial adult stage selected in mayflies involves extremely demanding conditions. However, mayflies have managed to overcome these by retaining an extra instar in the adult stage, thereby optimizing the water–air transition and facilitating the development required for flight, mating and reproduction.

## MATERIALS AND METHODS

A detailed description of the materials and methods is given in SI Appendix, Materials and Methods. In brief, the rearing methods of *C. dipterum* were as reported by Almudi et al. (14). RNA extraction and retrotranscription to cDNA and quantitative real-time PCR were performed according to the protocols described previously (18, 25). Primers used are detailed in *SI Appendix*, Table S1. For the statistical analyses of qRT-PCR measurements used the Relative Expression Software Tool (REST) (31). Methoprene treatments were carried out on freshly ecdysed last instar nymphs at a dose of 50 µg. For morphological studies and imaging we used a stereomicroscope Zeiss DiscoveryV8 and a bright field microscope Carl Zeiss-AXIO IMAGER.Z1.

## Supporting information

Supplemental Figures S1 and S2, supplemental Table S1, supplemental materrial and methods

## Data Availability

All study data are included in the main text and supporting information.

## Acknowledgements

Work supported by Spanish Ministry of Economy and Competitiveness (Grants CGL2015–64727-P, and PID2019-104483GB-I00 to XB), by Catalan Government (Grant 2017 SGR 1030 to XB), and by the European Fund for Economic and Regional Development (FEDER funds). OK received a Royal Thai Government Scholarship to do a PhD thesis in XB laboratory, in Barcelona, and AV-A, received a predoctoral fellowship of the Spanish Ministry of Economy and Competitiveness, associated to the Grant CGL2015–64727-P. IA was supported by the European Union’s Horizon 2020 research and innovation programme under the Marie Sklodowska-Curie Grant Agreement 657732.

## References

1. X. Belles, Insect metamorphosis. From natural history to regulation of development and evolution (Academic Press, 2020).

2. X. Belles, The innovation of the final moult and the origin of insect metamorphosis. Philos. Trans. R. Soc. B Biol. Sci. 374, 20180415 (2019).

3. J. Kukalová-Peck, Origin and evolution of insect wings and their relation to metamorphosis, as documented by the fossil record. J. Morphol. 156, 53–125 (1978).

4. J. Kukalová-Peck, Origin of the insect wing and wing articulation from the arthropodan leg. Can. J. Zool. 61, 1618–1669 (1983).

5. I. Almudi, et al., Genomic adaptations to aquatic and aerial life in mayflies and the origin of wings in insects. Nat. Commun. 11, 2631 (2020).

6. J. Lancaster, B. J. Downes, Aquatic Entomology (Oxford University Press, 2013).

7. R. L. Taylor, A. G. Richards, The subimaginal cuticle of the mayfly Callibaetis sp. (Ephemeroptera). Ann. Entomol. Soc. Am. 56, 418–426 (1963).

8. G. F. Edmunds, W. P. McCafferty, The mayfly subimago. Annu. Rev. Entomol. 33, 509–527 (1988).

9. V. C. Maiorana, Why do adult insects not moult? Biol. J. Linn. Soc. 11, 253–258 (1979).

10. X. Belles, C. G. Santos, The MEKRE93 (Methoprene tolerant-Krüppel homolog 1-E93) pathway in the regulation of insect metamorphosis, and the homology of the pupal stage. Insect Biochem. Mol. Biol. 52, 60–68 (2014).

11. R. E. Snodgrass, Insect metamorphosis. Smithson. Misc. Collect. 122, 1–124 (1954).

12. C. W. Schaefer, The mayfly subimago: a possible explanation. Ann. Entomol. Soc. Am. 68, 183 (1975).

13. F. P. Ide, The subimago of Ephoron leukon Will., and a discussion of the imago instar (Ephem.). Can. Entomol. 69, 25–29 (1937).

14. I. Almudi, et al., Establishment of the mayfly Cloeon dipterum as a new model system to investigate insect evolution. Evodevo 10, 6 (2019).

15. D. S. Brown, The morphology and function of the mouthparts of Cloeon dipterum L. and Baetis rhodani (Pictet) (Insecta, Ephemeroptera). Proc. Zool. Soc. Lond. 136, 147–176 (1963).

16. D. F. Erezyilmaz, L. M. Riddiford, J. W. Truman, The pupal specifier broad directs progressive morphogenesis in a direct-developing insect. Proc. Natl. Acad. Sci. U. S. A. 103, 6925–6930 (2006).

17. B. Konopová, M. Jindra, Broad-Complex acts downstream of Met in juvenile hormone signaling to coordinate primitive holometabolan metamorphosis. Development 135, 559–568 (2008).

18. J.-H. Huang, J. Lozano, X. Belles, Broad-complex functions in postembryonic development of the cockroach Blattella germanica shed new light on the evolution of insect metamorphosis. Biochim. Biophys. Acta -Gen. Subj. 1830, 2178–2187 (2013).

19. J. Cruz, D. Martín, X. Belles, Redundant ecdysis regulatory functions of three nuclear receptor HR3 isoforms in the direct-developing insect Blattella germanica. Mech. Dev. 124, 180–189 (2007).

20. R. J. Wootton, J. Kukalová-Peck, Flight adaptations in Palaeozoic Palaeoptera (Insecta). Biol. Rev. 75, 129–167 (2000).

21. T. Soldán, Secondary sexual characters in mayfly larvae and their evolutionary significance (Ephemeroptera). Acta ent. bohernoslov. 78, 140–142 (1981).

22. B. Konopová, V. Smykal, M. Jindra, Common and distinct roles of juvenile hormone signaling genes in metamorphosis of holometabolous and hemimetabolous insects. PLoS One 6, e28728 (2011).

23. M.-D. Piulachs, V. Pagone, X. Belles, Key roles of the Broad-Complex gene in insect embryogenesis. Insect Biochem. Mol. Biol. 40, 468–475 (2010).

24. P. R. DiBello, D. A. Withers, C. A. Bayer, J. W. Fristrom, G. M. Guild, The Drosophila broad-complex encodes a family of related proteins containing zinc fingers. Genetics 129, 385–397 (1991).

25. J. Lozano, X. Belles, Conserved repressive function of Krüppel homolog 1 on insect metamorphosis in hemimetabolous and holometabolous species. Sci. Rep. 1, 163 (2011).

26. E. Ureña, C. Manjón, X. Franch-Marro, D. Martín, Transcription factor E93 specifies adult metamorphosis in hemimetabolous and holometabolous insects. Proc. Natl. Acad. Sci. U. S. A. 111, 7024–7029 (2014).

27. Q. Si, J.-Y. Luo, Z. Hu, W. Zhang, C.-F. Zhou, De novo transcriptome of the mayfly Cloeon viridulum and transcriptional signatures of Prometabola. PLoS One 12, e0179083 (2017).

28. A. Berlese, Intorno alle metamorfosi degli insetti. Redia 9, 121–136 (1913).

29. H. Weber, Grundriss der Insektenkunde (Gustav Fischer, 1949).

30. J. Monod, Le hasard et la nécessite?: Essai sur la philosophie naturelle de la biologie moderne (Editions du Seuil, 1970).

31. M. W. Pfaffl, G. W. Horgan, L. Dempfle, Relative expression software tool (REST) for group-wise comparison and statistical analysis of relative expression results in real-time PCR. Nucleic Acids Res. 30, e36 (2002).

