## Supplemental Figures S1 and S2, supplemental Table S1, supplemental materrial and methods for "The mayfly subimago explained. The regulation of metamorphosis in Ephemeroptera"

SUPPORTING INFORMATION

**Material and Methods**

**Figure S1**. Alignment of the Broad complex Zinc fingers of *Cloeon dipterum*, with those of *Blattella germanica*, *Drosophila melanogaster*, *Ephemera danica* and *Ischnura elegans.*

**Figure S2.** Phylogenetic relationships of the Broad complex Zinc fingers of *Cloeon* dipterum, *Blattella germanica*, *Drosophila melanogaster*, *Ephemera danica* and *Ischnura elegans*.

**Table S1.** Primers used to measure expression levels of selected *Cloeon dipterum* genes by qRT-PCR.

**References cited**

### MATERIALS AND METHODS

#### *Cloeon dipterum* rearing in the laboratory

*C. dipterum* mayflies used in the experiments were obtained from a colony starting from gravid females by forced mating in the laboratory at Centro Andaluz de Biología del Desarrollo (Sevilla, Spain). The newly hatched nymphs were reared in unchlorinated and oxygenated water at 22 ± 1ºC, and under 12:12 h (light: dark) photoperiod, feeding them on filamentous algae (*Chara* sp.). A detailed description of the rearing methods through the entire life cycle is provided by Almudi et al. (1).

#### RNA extraction and retrotranscription to cDNA

Total RNA was extracted from whole bodies, body parts, or particular tissues of *C. dipterum* nymphs, subimago and adult, using the Gen Elute Mammalian Total RNA kit (Sigma-Aldrich), according to the manufacturer's instructions. RNA quantity and quality were estimated by spectrophotometric absorption at 260 nm in a Nanodrop Spectrophotometer ND-1000® (NanoDrop Technologies). A sample of 400 ng from each RNA extraction was then treated with DNase (Promega) and reverse transcribed with first Strand cDNA Synthesis Kit (Roche) and random hexamer primers (Roche).

#### Determination of mRNA levels by quantitative real-time PCR

Measurements with qRT-PCR were carried out in an iQ5 Real-Time PCR Detection System (Bio-Lab Laboratories), using SYBR®Green (iTaq™ Universal SYBR® Green Supermix; Applied Biosystems). Reactions were carried out in triplicate, and a template-free control was included in all batches. Primers used to measure the transcripts of interest are detailed in Table S1. The efficiency of each set of primers was validated by constructing a standard curve through three serial dilutions. Levels of mRNA were quantified relative to CdActin-5c mRNA (Table S1). Results are given as copies of the examined mRNA per 1000 copies of CdActin-5c mRNA.

**Statistical analyses of qRT-PCR measurements**

In all qRT-PCR experiments, to test the statistical significance differences between control and treated samples it has been used the Relative Expression Software Tool (REST), which evaluates the significance of the derived results by Pair-wise Fixed Reallocation Randomization Test (2).

**Treatments with methoprene**

To study the effect of juvenile hormone on metamorphosis, freshly ecdysed last instar nymphs of *C. dipterum* (less than 5 h after shedding off the exuvia) were treated with methoprene (Sigma-Aldrich) at a dose of 50 µg. Nymphs were immobilized on ice, and a volume of 0.25 µl of an acetone solution of methoprene (200 µg/µl) was topically applied on the mesonotum with a 5 µl Hamilton microsyringe. Controls received 0.25 µl of acetone.

**Morphological studies and imaging**

The nymphs, subimago and adults were examinations and photographs using a stereomicroscope Zeiss DiscoveryV8 and a bright field microscope Carl Zeiss-AXIO IMAGER.Z1.

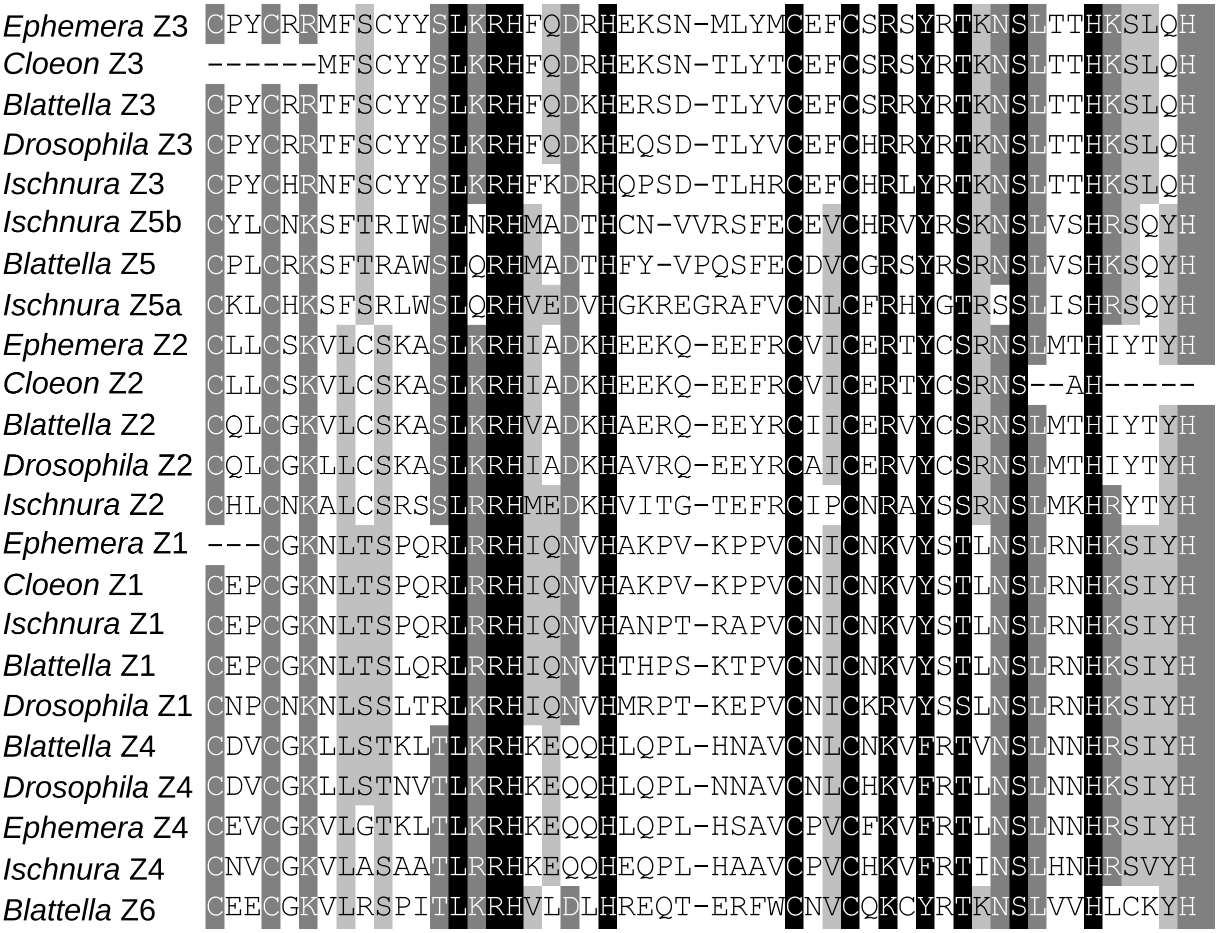


**Figure S1**. Alignment of the Broad complex Zinc fingers of *Cloeon dipterum*, with those of the neopteran especies *Blattella germanica* and *Drosophila melanogaster*, characterized by Piulachs et al. (3), and Di Bello et al. (4), respectively. Also included are those that we identified in the paleopteran species *Ephemera danica* and *Ischnura elegans*.The accession numbers of the Broad complex Zinc finger isoforms of *B. germanica* are: FN651774 (Z1), FN651775 (Z2), FN651776 (Z3), FN651777 (Z4), FN651778 (Z5), and FN651779 (Z6). Those of *D. melanogaster* are: CAA38474 (Z1), CAA38476 (Z2), CAA38475 (Z3), and AAB09760 (Z4). The Broad complex Zinc fingers of *C. dipterum*, *E. danica* and *I. elegans* were manually annotated from the respective genome projects PRJEB34721, PRJNA171755 and PRJNA353476. The alignment was carried out with Clustal X (5).

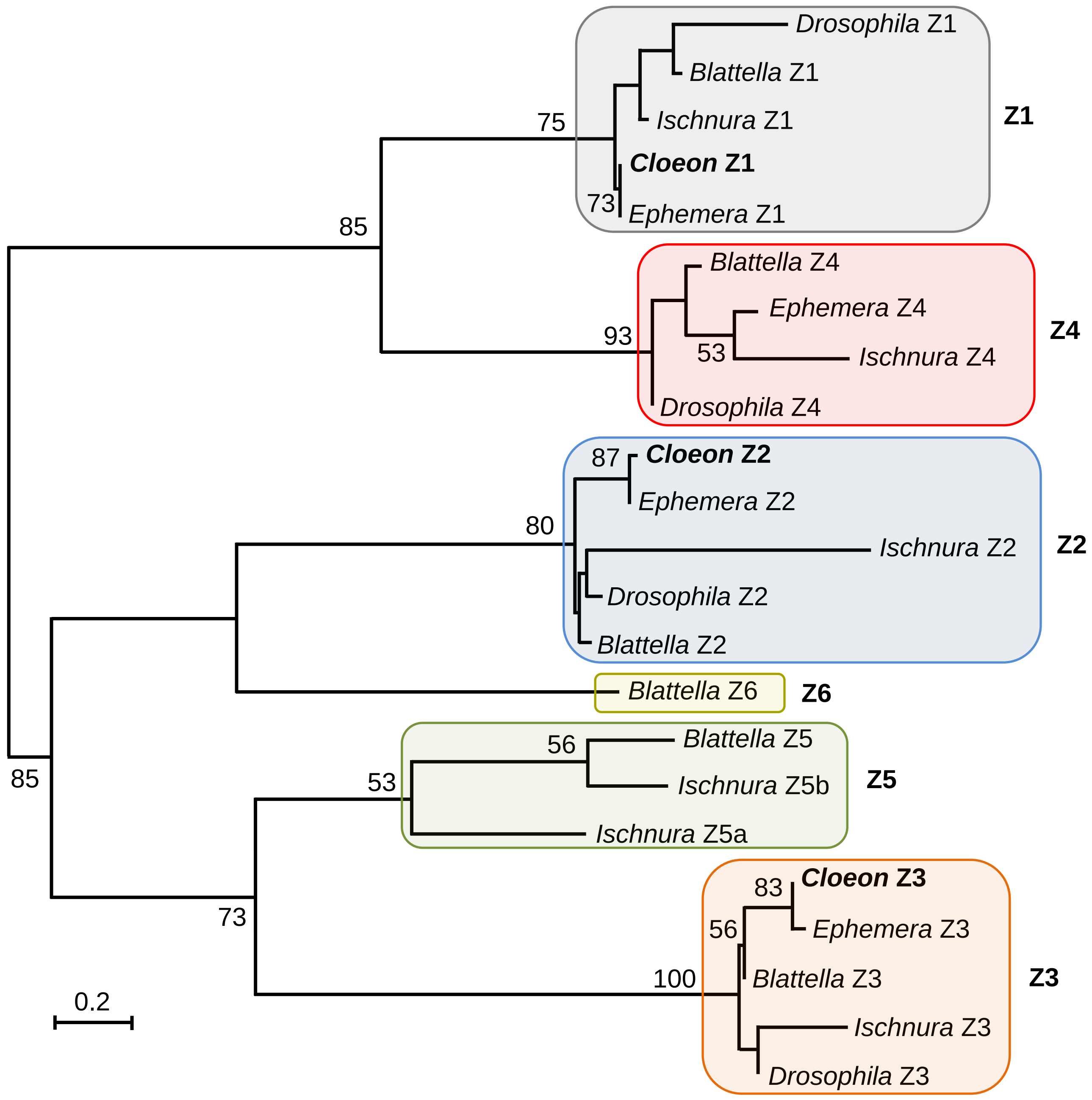


**Figure S2.** Phylogenetic relationships of the Broad complex Zinc fingers of *Cloeon dipterum*, *Blattella germanica*, *Drosophila melanogaster*, *Ephemera danica* and *Ischnura elegans*, according to the alignment and data showed in Figure S1. The phylogenetic reconstruction was carried out with PHYML 3.0 (6) based on the maximum-likelihood principle with the amino acid substitution model, four substitution rate categories, and a gamma shape parameter of 1.444. The data was bootstrapped for 100 replicates. Bootstrap values higher than 50 are indicated in the corresponding nodes. The sequences corresponding to *Cloeon dipterum* are indicated in bold. Scale bar indicates the number of substitutions per site.

**Table S1.** Primers used to measure expression levels of selected *Cloeon dipterum* genes by qRT-PCR

| Gene | Forward primer | Reverse primer |
|---|---|---|
| *Actin 5C* (*Act5C*) | AGAAGTTGCTGCCCTCGTT | GACCATCACACCCTGATGC |
| *Broad complex (Br-C core)* | AGGACTTCGTGGATGTGACC | TGCACATGGGGTACTCTTGA |
| *Broad complex Z1 (Br-C Z1* | GCTTGGTTATCGATGCGAAC | GTTTCGCGTGCACGTTCT |
| *Broad complex Z2 (Br-C Z2)* | TCGAAGGCTAGTCTCAAACGA | CGTTCGCAAATGACACATCT |
| *Broad complex Z3 (Br-C Z3)* | CTGTCGACGCATGTTCAGTT | CGTGTACAGCGTGTTGGACT |
| Ecdysone-induced protein 93F *(Eip93F, E93 or mblk-1)* | CTACGATCGTGACAGCCTGA | CGCTCCTTGACTTTGTACTCG |
| *Hormone receptor 3 (HR3)* | GCAACAAGAACTGCGTCGT | GCTGCTTTTTGGACATTCGT |
| *Krüppel homolog 1* (*Kr-h1*) | TGCGAGTACTGCCACAAGTC | CATTTGTACGGTCGCTCCTT |

Note: Genes manually annotated in *Cloeon dipterum* genome project accessions PRJEB34721. The sequence reads and the genome assembly have been deposited in the European Nucleotide Archive (ENA).

### References cited in the supporting information


1. Almudi I, Martín-Blanco CA, García-Fernandez IM, López-Catalina A, Davie K, Aerts S, Casares F (2019) Establishment of the mayfly *Cloeon dipterum* as a new model system to investigate insect evolution. *Evodevo* 10 (1): 6.

2. Pfaffl MW, Horgan GW, Dempfle L (2002) Relative expression software tool (REST) for group-wise comparison and statistical analysis of relative expression results in real-time PCR. *Nucleic Acids Res* 30 (9): e36.

3. Piulachs M-D, Pagone V, Belles X (2010) Key roles of the Broad-Complex gene in insect embryogenesis. *Insect Biochem Mol Biol* 40 (6): 468–475.

4. DiBello PR, Withers DA, Bayer CA, Fristrom JW, Guild GM (1991) The *Drosophila* broad-complex encodes a family of related proteins containing zinc fingers. *Genetics* 129: 385–397.

5. Larkin MA, Blackshields G, Brown NP, Chenna R, McGettigan PA, McWilliam H, Valentin F, Wallace IM, Wilm A, Lopez R, Thompson JD, Gibson TJ, Higgins DG (2007) Clustal W and Clustal X version 2.0. *Bioinformatics* 23 (21): 2947–2948.

6. Guindon S, Gascuel O (2003) A simple, fast and accurate algorithm to estimate large phylogenies by maximum likelihood. *Syst Biol* 52: 696–704.
